# Mechanical analysis of spatiotemporal traction stress dynamics in a bleb-driven migrating cell, *Amoeba proteus*

**DOI:** 10.64898/2026.06.11.728063

**Authors:** Rio Terauchi, Syun Echigoya, Charles Fosseprez, Atsushi Taniguchi, Takuya Ohmura, Jean-Paul Rieu, Katsuhiko Sato, Toshiyuki Nakagaki, Yukinori Nishigami

**Affiliations:** Graduate School of Life Science, Hokkaido University, Sapporo, Japan; Research Institute for Electronic Science, Hokkaido University, Sapporo, Japan; Institute for Integrated Innovations, Hokkaido University, Sapporo, Japan; Department of Biomedical Engineering, Tohoku University, Sendai, Japan; Institut Lumière Matière, Université Lyon 1, CNRS, UMR5306, Villeurbanne, France; Program of Mathematics and Informatics, Faculty of Science, University of Toyama, Toyama, Japan

**Keywords:** amoeboid locomotion, bleb-driven locomotion, *Amoeba proteus*, traction stress, biophysical mechanics

## Abstract

Many adherent eukaryotic cells exhibit amoeboid locomotion, where traction stress exerted on the substrate is essential for movement. In this study, we investigated the spatiotemporal development of these forces in *Amoeba proteus* to clarify the mechanical dynamics underlying bleb-driven migration. By performing a multipole analysis of the stress distribution, we characterized the spatiotemporal patterns exhibited by motile cells. Furthermore, we tracked the behavior of individual localized peak structures within these profiles, which are thought to correspond to focal contact sites. These analyses revealed that the front-back asymmetry in the traction distribution correlates with the direction of migration. We also found that *A. proteus* exhibits a periodic pattern in which inward-directed stresses are alternately strengthened and weakened at the cell poles. Crucially, we identified a distinctive feature not observed in other cell types: the generation of large lateral traction forces at the cell center. Together, these results highlight both the universality and diversity of the biophysical mechanisms driving amoeboid locomotion.

## Introduction

Amoeboid locomotion is widely observed across adherent eukaryotic cells, ranging from unicellular to multicellular organisms. This type of locomotion is involved in a wide range of biological processes, including wound healing[1–3], development[4–7], cancer metastasis[8,9], and immune responses[10,11]. In addition, amoeboid locomotion is observed in a variety of protists[12–14]. The mechanisms underlying this motility are broadly classified into two modes: bleb-driven and actin polymerization-driven migration[15–19]. In the latter, propulsion is generated by actin polymerization at the leading edge of pseudopodia, whereas bleb-driven locomotion relies on contractile forces generated by cortical actomyosin[15,16,20–22]. Moreover, some cell types can switch between these modes depending on their internal state and the surrounding environment[17,20–23]. Actin polymerization-driven locomotion is mainly observed in strongly adhesive two-dimensional (2D) environments[17,23], or in less dense three-dimensional (3D) matrices[16,24]. In contrast, bleb-driven locomotion is typically observed under weak-adhesion conditions^15,17–19^ or within mechanically confined 3D environments, such as tissues[15–19].

To achieve amoeboid locomotion, cells must exert stress on their external environment[16,18,19,25]. This interaction, generally referred to as traction stress, is essential for motility and has been measured on 2D substrates using traction force microscopy (TFM)[26–32]. In cells exhibiting actin polymerization-driven locomotion, these forces have been extensively characterized through traction force microscopy (TFM), which has revealed that traction stress distributions correlate with the direction of cell migration[29,31] and exhibit characteristic localized regions of high magnitude[29,30], often termed traction hotspots.

While traction stresses have been extensively characterized in cells using actin polymerization-driven amoeboid locomotion in 2D, far less is known about their organization during bleb-driven amoeboid locomotion, a topic that remains a central focus of modern cell biology[19].This gap stems partly from the fact that blebbing motility often occurs under weak adhesion or within confined 3D environments, where traction force measurements are technically challenging. The free-living *Amoeba proteus* (*A. proteus*) provides an ideal model system to address this question, as it exhibits bleb-driven locomotion even on 2D substrates[19,33,34], where TFM is well established. In this study, we investigated the spatiotemporal development of traction forces in *A. proteus* to clarify the mechanisms of bleb-driven amoeboid locomotion.

## Results

### a. Stress field and amoeboid locomotion

*A. proteus* forms adhesive contacts on the substrate and exerts traction stress during amoeboid locomotion. To elucidate the mechanical basis of amoeboid locomotion in *A. proteus*, we measured the traction stress ***F*** during migration (Fig. 1A; Supplementary Movie S1 and S2). The time- and sample-averaged traction stress vectors were predominantly directed toward the cell centroid (Fig. 1B). When decomposed into components parallel and perpendicular to the migration direction, the parallel component 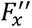 exhibited a bipolar spatial distribution, whereas the perpendicular component 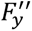 showed enhanced magnitudes near the cell center (Fig. 1C).

**Fig. 1.**
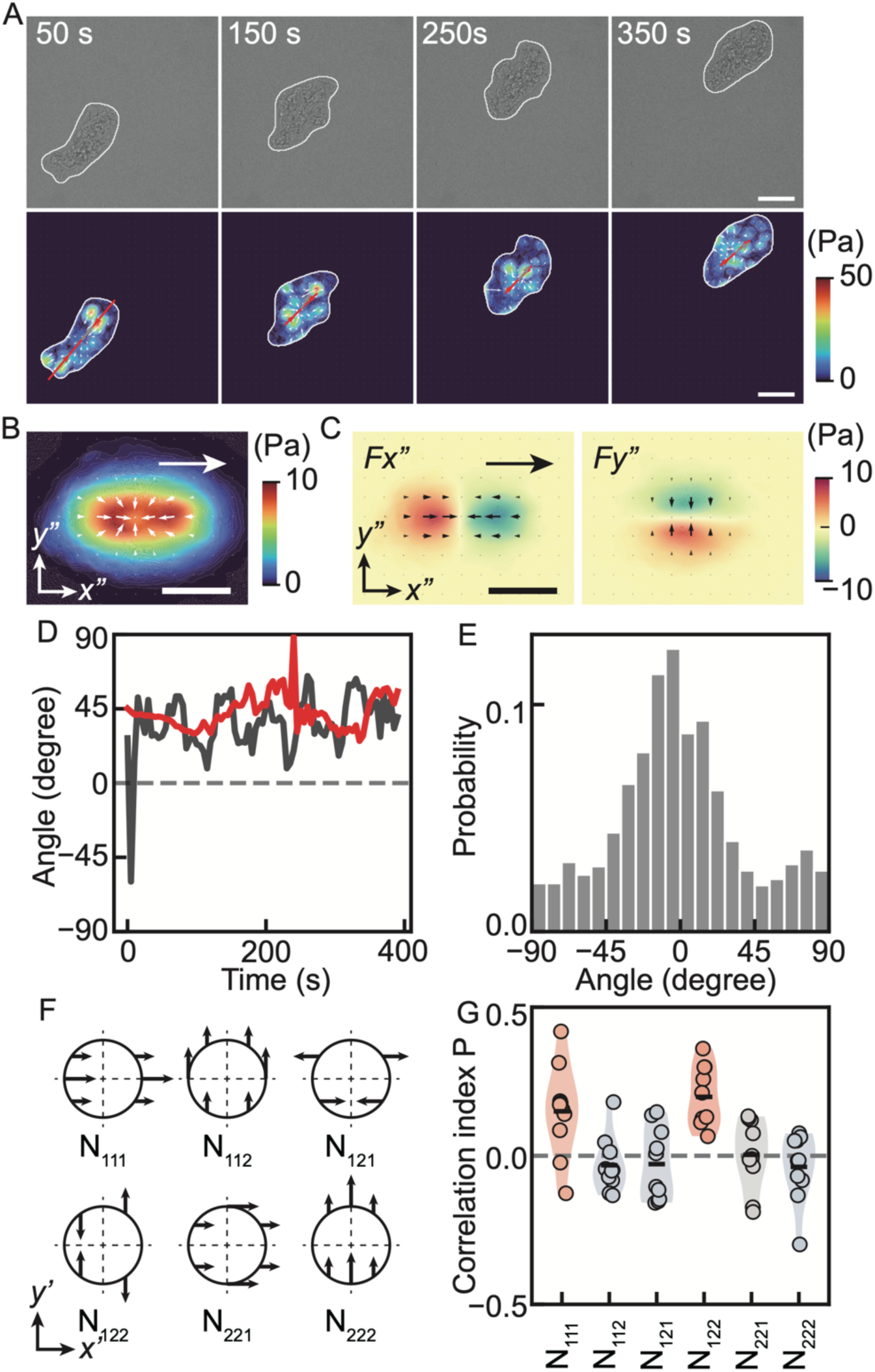
Spatiotemporal traction stress distribution and multipole analysis during cell migration. (A) Time evolution of cell migration and traction stress distribution. (Top) Bright-field images of migrating *A. proteus* after background subtraction. White contours indicate the cell boundary. (Bottom) Corresponding traction stress fields during migration. The color bar indicates stress magnitude. Red arrows denote the major dipole, and gray arrows indicate the cell velocity vector. Scale bars, 100 µm. (B) Time- and cell-averaged traction stress distribution. The analysis was performed in a reference frame fixed to the cell centroid and migration direction (white arrow). Scale bar, 100 µm. N = 10 cells. (C) Decomposition of the averaged traction stress distribution into components parallel (*Fx*″, left) and perpendicular (*Fy*″, right) to the direction of migration. Scale bar, 100 µm. N = 10 cells. (D) Temporal evolution of the angle between the cell velocity vector and the major dipole in the experimental reference frame. (E) Probability distribution function (PDF) of the angle between the velocity vector and the major dipole. N = 10 cells. (F) Six components of the second-moment tensor. The indices *i*, *j*, *k* correspond to *x*_*i*_, *y*_*j*_, and 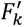 in Eq. [8], where each index takes values 1 or 2 corresponding to the x and y components, respectively. (G) Violin plot of the correlation index *P* between the cell velocity vector and the second-moment tensor. Positive values indicate a tendency for aligned signs, negative values indicate anti-correlation, and zero indicates no correlation. Dots represent individual cells. Black lines and violin colors indicate the mean value of the correlation index.

To quantify the relationship between traction stress organization and amoeboid locomotion, we performed a multipole analysis²⁹. We first computed the first-order moment (force dipole) *M*_*ij*_ (Eq. [8] in Materials and Methods). Diagonalization of *M*_*ij*_ yielded two eigenvalues and corresponding eigenvectors. The eigenvector associated with the larger (smaller) absolute eigenvalue is defined as the major (minor) dipole, and the direction of the major dipole is referred to as the dipole axis. The velocity vector was found to be aligned with the dipole axis throughout the observation period (Fig. 1D). This alignment was quantified by computing the probability density function (PDF) of the angle between the velocity vector and the dipole axis, which exhibited a peak around 0° (Fig. 1E). This indicates a preferred alignment between the migration direction and the force dipole axis, although it does not specify the directionality of migration.

To address this, we analyzed the second-order moment (force quadrupole) *N*_*ijk*_ (Eq. [9] in Materials and Methods), which consists of six independent components (Fig. 1F). We quantified the consistency between the sign of each quadrupole component and the sign of the velocity component along the migration direction (Fig. 1G). In particular, the components *N*_111_ and *N*_122_ showed strong positive correlations with the direction of migration. The positive correlation in *N*_111_ and *N*_122_ indicates that traction stress components parallel and perpendicular to the migration direction are biased toward the back of the cell.

### b. Traction stress dynamics during amoeboid locomotion

To investigate correlations in traction stress dynamics, we analyzed the spatiotemporal evolution of the traction stress. First, the migration direction of each cell was aligned so that all cells migrated in the same direction (Fig. 2A, B). We then integrated the traction stress components parallel and perpendicular to the migration direction, 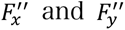, across the cell width to obtain kymographs of the integrated forces *F*_*tx*_ (Fig. 2C) and *F*_*ty*_ (Fig. 2D) (see Materials and Methods for details). Both kymographs exhibited band-like patterns parallel to the horizontal axis that periodically emerged from the cell front. We refer to these band-like structures as traction patterns. The fact that these patterns are approximately parallel to the horizontal axis indicates that the traction sources remain nearly stationary relative to the substrate while the cell migrates forward. In the cell reference frame, these traction patterns therefore move from the front toward the back of the cell.

**Fig. 2.**
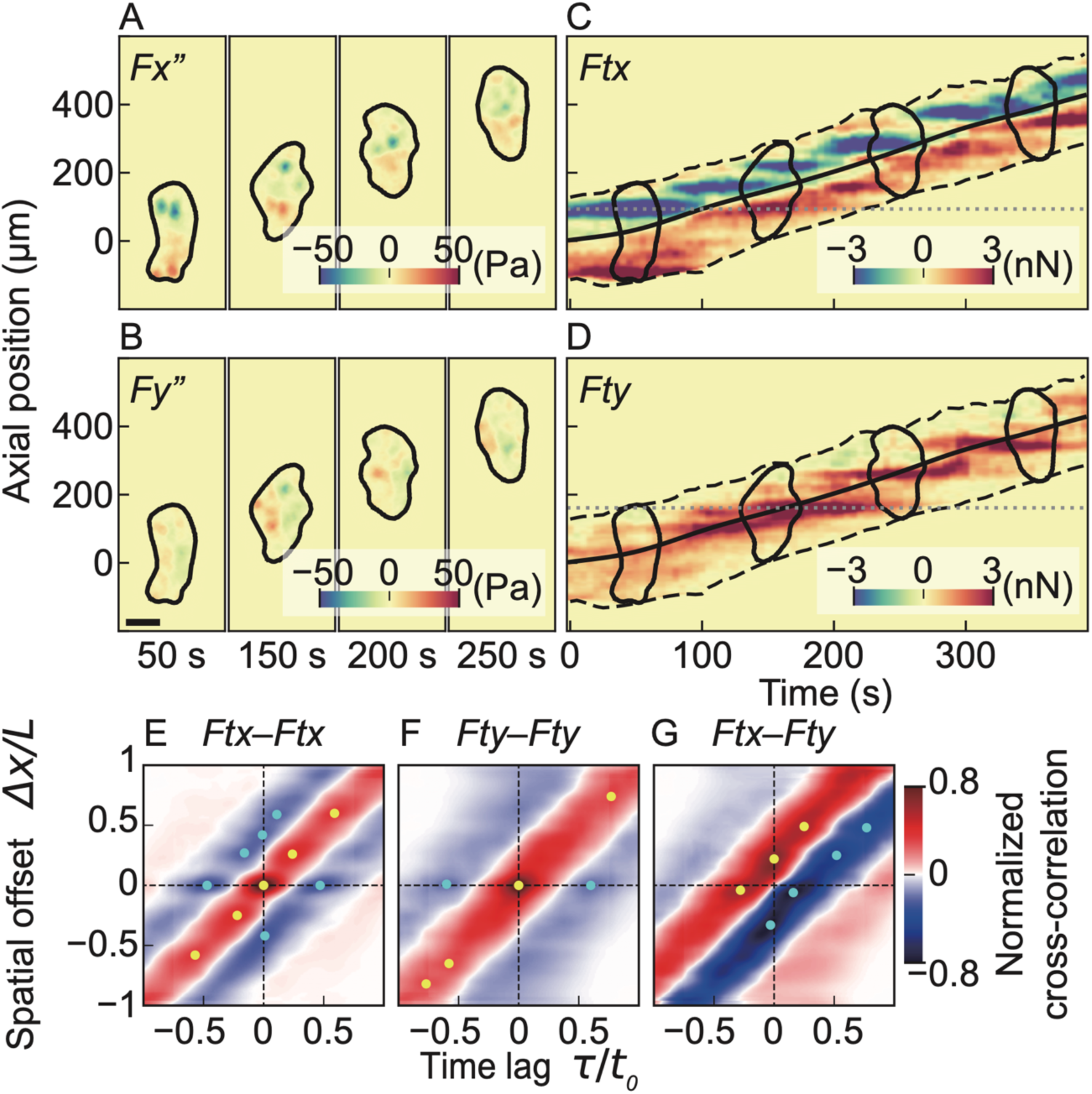
Spatiotemporal development of traction stress during amoeboid locomotion. (A) Time evolution of amoeboid locomotion and the traction stress component parallel to the migration direction 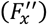, shown after aligning the migration direction upward. (B) Time evolution of amoeboid locomotion and the traction stress component perpendicular to the migration direction 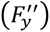. The origin of the axial position was defined as the cell centroid at *t* = 0. Cell contours are shown as black solid lines. Scale bar, 100 µm. (C) Temporal evolution of the integrated traction stress component parallel to the migration direction (*F*_*tx*_) along the migration axis. (D) Temporal evolution of the integrated perpendicular component (*F*_*ty*_) along the migration axis. In (C) and (D), each column represents the net traction force integrated across the cell width at each time point, expressed in nN. For the perpendicular component, inward-directed stresses were defined as positive to avoid cancellation across the cell width. Cell contours corresponding to the same time points as in (A) and (B) are shown. Black solid lines indicate the trajectory of the cell center, and black dashed lines indicate the trajectories of the front and back edges of the cell. Gray dotted lines indicate one of the reference axial positions used for the correlation analysis. (E) Two-dimensional fluctuation cross-correlation function (2D FCC) between the parallel traction force components (*F*_*tx*_–*F*_*tx*_). (F) 2D FCC between the perpendicular traction force components (*F*_*ty*_–*F*_*ty*_). (G) 2D FCC between the parallel and perpendicular traction force components (*F*_*tx*_–*F*_*ty*_). Spatial offset and time lag were normalized. A spatial offset of 1 corresponds to the mean cell length, and a time lag of 1 corresponds to the time required for a cell to migrate a distance equal to its own length. To compensate for inter-sample variability, each 2D FCC map was normalized by its maximum absolute value before averaging across samples. Yellow dots indicate detected positive peaks, and cyan dots indicate detected negative peaks. N = 10 cells.

Using these kymographs, we calculated RMS-weighted averaged two-dimensional fluctuation cross-correlation functions (2D FCCs; Eq. [11] in Materials and Methods) as functions of normalized time lag *τ*/*t*_0_ and normalized spatial offset Δ*x*/*L* for *F*_*tx*_-*F*_*tx*_ (Fig. 2E), *F*_*ty*_-*F*_*ty*_ (Fig. 2F), and *F*_*tx*_-*F*_*ty*_ (Fig. 2G), respectively (N = 10 cells). Here, the time lag was normalized by the time required for a cell to migrate one cell length, *t*_0_, and the spatial offset was normalized by the mean cell length, *L*. Thus, a normalized time lag of 1 corresponds to the time required for a cell to move its own length, and a normalized spatial offset of 1 corresponds to one mean cell length. All three 2D FCCs exhibited diagonal ridge-like structures approximately parallel to Δ*x*/*L* = *τ*/*t*_0_, along which positive and negative peaks appeared at roughly regular intervals (Fig. 2E–G). These patterns suggest that traction patterns repeatedly emerge at similar positions within the cell.

In the F_tx_ kymograph, from front to tail, traction patterns first appeared with negative values, subsequently switched to positive values, and then disappeared (Fig. 2C). This reflects the generation of backward-directed forces at the front of the cell and forward-directed forces at the back. In the 2D FCC of *F*_*tx*_-*F*_*tx*_, negative peaks were observed around Δ*x*/*L* = 0 at *τ*/*t*_0_ = ±0.47 (Fig. 2E). These peaks suggest that the backward and forward force peaks are separated by approximately 0.5 cell lengths. At individual time points in the *F*_*tx*_ kymograph, three traction patterns were often observed simultaneously within a single cell: a newly formed pattern near the front, an older pattern near the cell center, and another older pattern near the back (Fig. 2C, after 100 s). In the region −0.5 < Δ*x*/*L* < 0.5 and −0.5 < *τ*/*t*_0_ < 0.5, positive peaks were detected at (*τ*/*t*_0_, Δ*x*/*L*) = (−0.22, −0.25), (0, 0), and (0.24, 0.26)(Fig. 2E). These peaks suggest that neighboring traction patterns are spaced at intervals of approximately 0.25 cell lengths apart, consistent with the direct observations in the kymograph. Returning to the *F*_*tx*_kymograph, the traction patterns appeared sequentially in time. Consequently, when a newly formed traction pattern reached a strong negative peak, the older pattern near the cell center was close to zero, whereas an even older pattern near the back reached a strong positive peak (Fig. 2C). Consistent with this observation, the 2D FCC of *F*_*tx*_-*F*_*tx*_ showed negative peaks around *τ*/*t*_0_ = 0 at Δ*x*/*L* = −0.43 and 0.42 (Fig. 2E). These peaks indicate that strong backward-directed forces at the front coincide with strong forward-directed forces at the back.

In contrast, in the *F*_*ty*_kymograph, traction patterns gradually increased in magnitude, reached a peak near the cell center, and subsequently weakened (Fig. 2D). Although traction patterns also appeared periodically in *F*_*ty*_, the spacing between neighboring patterns exhibited greater variability than that observed for *F*_*tx*_ (Fig. 2D). In the 2D FCC of *F*_*ty*_-*F*_*ty*_, negative peaks were observed around Δ*x*/*L* = 0at *τ*/*t*_0_ = ±0.70 (Fig. 2F), suggesting the characteristic duration of the *F*_*ty*_traction patterns. On the other hand, in the region −0.5 < Δ*x*/*L* < 0.5 and −0.5 < *τ*/*t*_0_ < 0.5, positive peaks were not detected (Fig. 2F). This suggests that *F*_*ty*_ traction patterns emerge more randomly than *F*_*tx*_ traction patterns.

Finally, we examined the relationship between *F*_*tx*_ and *F*_*ty*_. The two traction patterns generally appeared at similar axial positions (Fig. 2C, D). In the 2D FCC of *F*_*tx*_-*F*_*ty*_, around Δ*x*/*L* = 0, a positive peak was observed at *τ*/*t*_0_ = −0.28, whereas a negative peak was observed at *τ*/*t*_0_ = 0.16 (Fig. 2G). This asymmetry likely reflects the front-back asymmetry of the traction stress distribution (see Discussion). Together, these results reveal characteristic spatiotemporal patterns in the traction forces generated by *A. proteus* during amoeboid locomotion.

### c. Detection and analysis of contact sites dynamics

*A. proteus* adheres to the substrate through patch-like contact regions and exerts traction stress on the substrate from these regions (Fig. 3A, left column). We refer to these regions as contact sites. These contact sites underlie the spatial distribution and spatiotemporal dynamics of traction stress described in the previous sections. Here, we focused on the dynamics of individual contact sites and investigated their contributions to the macroscopic spatiotemporal features of the traction stress.

**Fig. 3.**
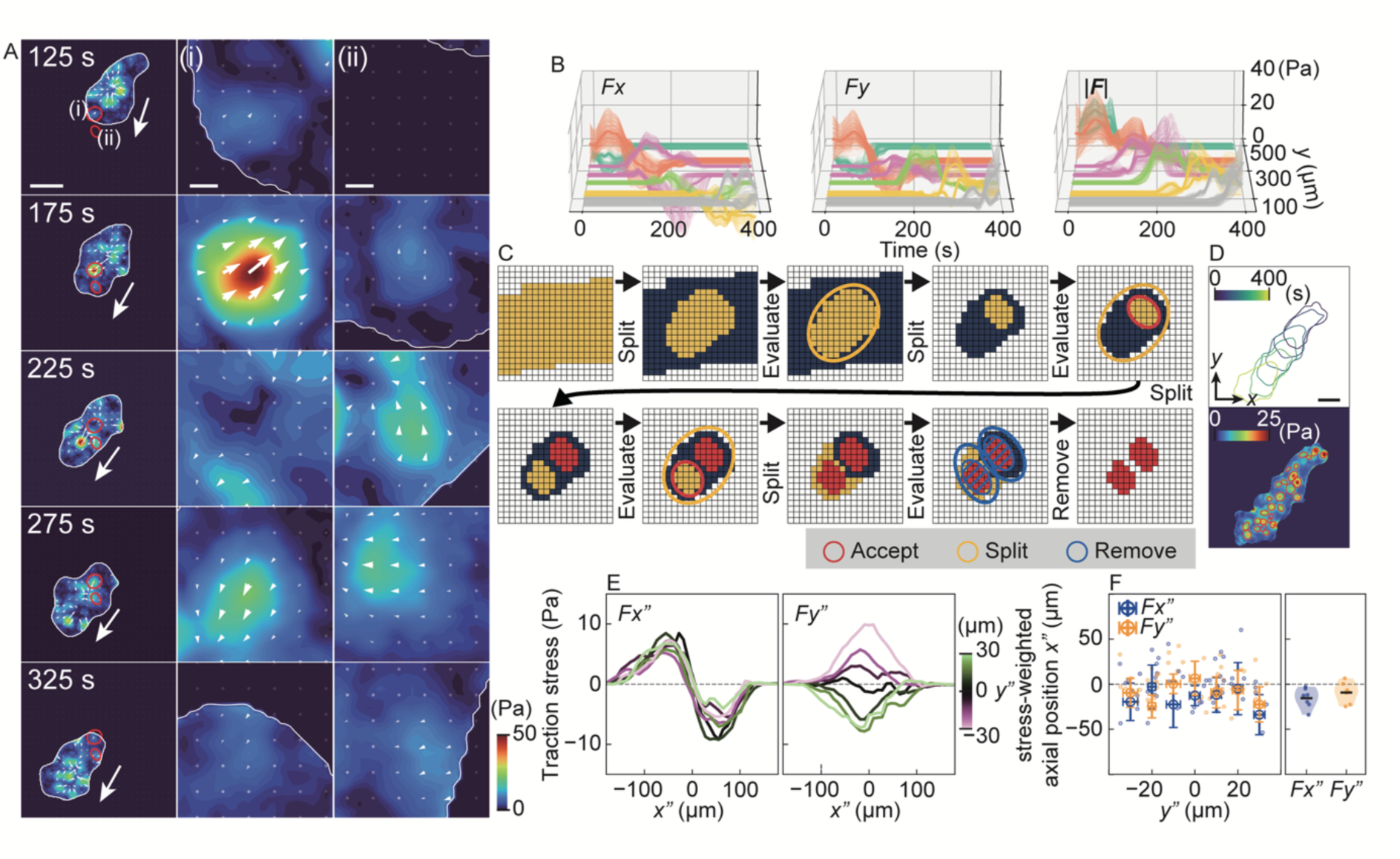
Dynamics of individual contact sites. (A) Dynamics of contact sites during cell migration. (Left column) Cell migration and contact sites observed in a wide-field view. Two representative contact sites are highlighted by red ellipses (i, ii). White arrows indicate the migration direction. Scale bar, 100 µm. (Center column) Magnified view of the dynamics of contact site (i), located near the cell center axis. Scale bar, 10 µm. (Right column) Magnified view of the dynamics of contact site (ii), located away from the cell center axis. Scale bar, 10 µm. In the magnified views, the image center was aligned to the center of each contact site. (B) Temporal evolution of traction stress generated by pixels classified into each contact-site cluster. Curve colors correspond to different contact sites. Semi-transparent curves indicate the temporal evolution of traction stress at individual pixels, whereas thick curves indicate the cluster-averaged temporal dynamics. The horizontal axis represents time (*t*), the depth axis represents the *y*-coordinate (µm), and the vertical axis represents traction stress magnitude (Pa). (C) Schematic of contact-site cluster detection. Yellow and dark blue regions indicate those segmented by the algorithm, whereas red regions indicate those that satisfy the criteria for contact sites. Blue ellipses indicate ellipses fitted to regions that did not satisfy the contact-site criteria, whereas red ellipses indicate ellipses fitted to regions satisfying the criteria. (D) Detected contact sites. (Top) Cell trajectories color-coded by time. Scale bar, 100 µm. (Bottom) Time-averaged traction stress distribution and detected contact sites (red ellipses). The time averaging was normalized at each pixel by the number of frames in which that pixel was included within the cell mask. (E) Relationship between the position along the cell center axis and traction stress generated by contact sites. (Left) Component parallel to the migration direction. (Right) Component perpendicular to the migration direction. Curve colors indicate the position *y*″(µm) relative to the cell center axis. (F) Stress-weighted axial position *x*″ of the traction stress distribution. The stress-weighted axial position *x*″ was defined as the weighted average of the coordinate *x*″, with the weights given by the absolute values of the traction stresses. (Left) Dependence of the stress-weighted axial position *x*″ of the traction stress distribution on the lateral position *y*″. Small open circles indicate values obtained from individual contact sites, whereas large open circles represent values obtained from representative stress distribution in each *y*″ bin. Error bars represent the ± standard deviation within each *y*″ bin. Blue and orange symbols denote the traction stress components 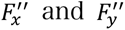, respectively. (Right) Violin plot of stress-weighted axial position *x*″ obtained from the representative traction stress distributions of all *y*″ bins.

Once formed, contact sites remained approximately fixed relative to the substrate and continuously exerted traction stress directed toward the cell center until detachment. The behavior of the contact sites depended on their position relative to the cell centerline. The contact site located near the cell centerline (i) initially strengthened the backward traction stress at the front of the cell, weakened the stress as it approached the cell center, subsequently exerted forward traction stress as it moved toward the back, and eventually detached from the substrate (Fig. 3A, center column). In contrast, the contact site located away from the centerline (ii) exerted weaker traction stress parallel to the migration direction but showed a larger traction stress component perpendicular to the migration direction (Fig. 3A, right column).

To extract these contact sites from the two-dimensional stress field, we analyzed the temporal evolution of the traction stress magnitude |*F*| and its *x*- and *y*-components (*F*_*x*_, *F*_*y*_) in the fixed laboratory frame (Fig. 3B). Contact sites were identified as groups of neighboring pixels exhibiting similar temporal dynamics using k-means clustering[35,36] (Fig. 3C; see Materials and Methods for details). Using this procedure, we successfully extracted the contact sites formed by *A. proteus* (Fig. 3D).

We next investigated the relationship between the axial position along the cell centerline and the traction stress generated at contact sites (Fig. 3E). The traction stress component parallel to the migration direction was typically directed backward (negative) at the front of the cell, weakened near the cell center, and became forward-directed (positive) at the back. In contrast, the traction stress component perpendicular to the migration direction increased near the cell center, and contact sites farther from the cell centerline tended to exert larger perpendicular traction stress (Fig. 3E, right). Taken together, these results suggest contact sites exhibit position-dependent dynamics while consistently generating traction stress directed toward the cell center.

We also found that both traction stress components parallel and perpendicular to the migration direction, 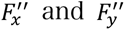, were slightly biased toward the back of the cell (Fig. 3E). To quantify this rearward bias, we calculated the stress-weighted axial position of the traction distribution along the migration axis *x*″. Specifically, the value was defined as the weighted average of the coordinate along the cell centerline, *x*″, using the absolute magnitude of each traction stress component as the weight. We then examined how this position depended on the lateral position, *y*″, of the contact site (Fig. 3F, left). The stress-weighted axial position obtained from representative 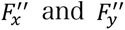 distributions were -15.4 ± 10.8 µm and -9.30 ± 11.3 µm (mean ± SD, N = 7), respectively (Fig. 3F, right). These results indicate that the traction stresses generated at contact sites exhibit front-back asymmetry, with their distributions biased toward the back of the cell.

## Discussion

We investigated the relationship between traction stress distribution and amoeboid motility in *A. proteus*, a cell that exhibits bleb-driven locomotion with TFM. In this study, we focused on contact sites formed by cells on the substrate and analyzed their individual dynamics and the spatiotemporal structure of the resulting traction stress distribution. Through these analyses, we investigated how the dynamics of traction stress generated at contact sites in *A. proteus* contribute to the amoeboid motility.

We first characterized the average spatial structure of the traction stress distribution. By averaging the traction stress over time and across cells, we found that stresses at the front of the cell were directed opposite to the direction of migration, whereas those at the back were directed along the migration direction, indicating that traction stresses were generally oriented toward the cell centroid (Fig. 1B). This centripetal organization is a well-established hallmark of cell-substrate interactions for both fibroblast or fast amoeboid cells like *Dictyostelium*, reflecting the inward pull generated by the contractile actomyosin cortex to maintain mechanical equilibrium[37,38]. We then decomposed the traction stress into components parallel and perpendicular to the direction of cell migration (Fig. 1C).

The parallel component was concentrated at both the front and back of the cell, whereas the perpendicular component was predominantly localized in the central region of the cell. It should be noted that the magnitudes in the averaged traction maps represent time- and sample-averaged stresses over the entire cell region. Because traction generation in *A. proteus* is spatially localized to discrete contact sites, many regions of the cell-substrate interface exert little traction at a given time. Averaging over these low-stress regions and over contact sites that appear at different positions across different cells and time points therefore reduces the apparent magnitude of the averaged stress field. Thus, the averaged maps should be interpreted primarily as showing the spatial organization of traction stresses rather than the peak stresses generated at individual contact sites.

We next performed a multipole analysis[29] to further assess the relationship between the traction stress distribution and the migration direction. With the first moment of the stress field, we found that the dipole axis aligns with the migration axis (Fig. 1D, E). We then analyzed the asymmetry of the stress field to relate it to the migration direction, using six components of the force quadrupole (Fig. 1F). As a result, we found *N*_111_ and *N*_122_ had a positive correlation with the cell velocity parallel to the dipole axis (Fig. 1G). The positive correlation in *N*_111_ and *N*_122_indicates that traction stress components parallel and perpendicular to the migration direction are biased toward the back of the cell. This result suggests that the front-back asymmetry of the traction stress field is correlated with the migration direction.

Such asymmetry in the traction stress distribution and its relationship with the direction of migration has also been observed not only in bleb-driven amoeboid locomotion but also in other cells, including cells that exhibit actin polymerization-driven amoeboid locomotion, another major mode of amoeboid migration[29,31,39]. In general, the traction stress generated by a cell balances internally, resulting in a net stress of zero[29]. Such kinematic constraints, independent of locomotion mode, may contribute to the conserved relationship between traction stress asymmetry and migration direction observed across species and motility modes.

We next examined the spatiotemporal patterns of the traction stress distribution using two-dimensional cross-correlation analysis. The cell migration direction was fixed, and the traction stress was decomposed into components parallel and perpendicular to migration (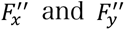), which were integrated across the cell width to obtain *F*_*tx*_ and *F*_*ty*_, respectively. The temporal evolution of these quantities along the migration axis was then visualized as kymographs (Fig. 2C, D). In these kymographs, band-like patterns (traction patterns) parallel to the horizontal axis appeared repeatedly from the leading edge. To quantitatively characterize the relationships between these patterns, we performed two-dimensional cross-correlation analyses in time-space coordinates (temporal lag and spatial shift) (Fig. 2E–G).

For *F*_*tx*_, the distribution of peaks in the two-dimensional cross-correlation map (Fig. 2E) indicated a characteristic spatial periodicity of approximately 0.25 cell lengths. Although traction-force patterns in *F*_*ty*_ also appeared with a comparable spacing, the corresponding cross-correlation map did not exhibit a clear peak at this interval (Fig. 2F). This is likely because the periodic appearance of *F*_*ty*_ patterns was less regular, owing to intermittent disappearance or missing events in the pattern sequence (e.g., around 50 s in Fig. 2D). As a result, variability in the spacing between successive patterns reduced the coherence of the correlation signal, causing the expected peak to become indistinct in the two-dimensional cross-correlation map. In the cross-correlation between *F*_*tx*_ and *F*_*ty*_ (Fig. 2G), a positive correlation peak was observed at Δ*x*/*L* ≈ 0 and *τ*/*t*_0_ = −0.28, while a negative correlation peak was found at *τ*/*t*_0_ = 0.16. Given that *F*_*ty*_reaches its maximum near the central region of the cell, these results indicate that backward-directed traction stresses peak approximately 0.16 cell lengths ahead of the cell center, whereas forward-directed traction stresses peak approximately 0.28 cell lengths behind it. These results suggest that traction stress generation is spatially asymmetric with respect to the direction of migration, with a bias toward the back of the cell. This is consistent with the analysis based on the quadrupole *N*_111_ (Fig. 1F, G).

The spatial features and spatiotemporal patterns of traction stress described above originate from patch-like adhesion regions formed on the substrate by the cell (Fig. 1A, bottom; Fig. 3A; Supplementary Movie S2). Focusing on these contact sites, we identified individual regions directly from the stress field patterns (Fig. 3B–D) and further analyzed their dynamics from formation to detachment. Our analysis revealed a common dynamic behavior: contact sites form at the leading edge, generate traction stresses consistently directed toward the cell centroid, and subsequently translocate toward the back of the cell, where they eventually detach (Fig. 3A; Supplementary Movie S2). The traction stress generated by individual contact sites exhibited distinct spatial characteristics. Focusing on the traction stress component parallel to the migration direction, we found that contact sites at the front of the cell exerted backward-directed stresses, whereas those at the back exerted forward-directed stresses (Fig. 3E, left; Fig. 4A, left). We next examined the perpendicular component of traction stress and found that its magnitude tended to increase with increasing distance of the contact site from the cell centerline (Fig. 3E, right; Fig. 4A, right). To further quantify the spatial bias of traction generation, we calculated the stress-weighted axial position of the traction stress distribution along the cell centerline. For both the parallel and perpendicular components, the position was located slightly behind the cell center (Fig. 3F). These findings suggest that the front-back asymmetric behavior of contact sites contributes to the front-back asymmetry of the overall traction stress distribution.

**Fig. 4.**
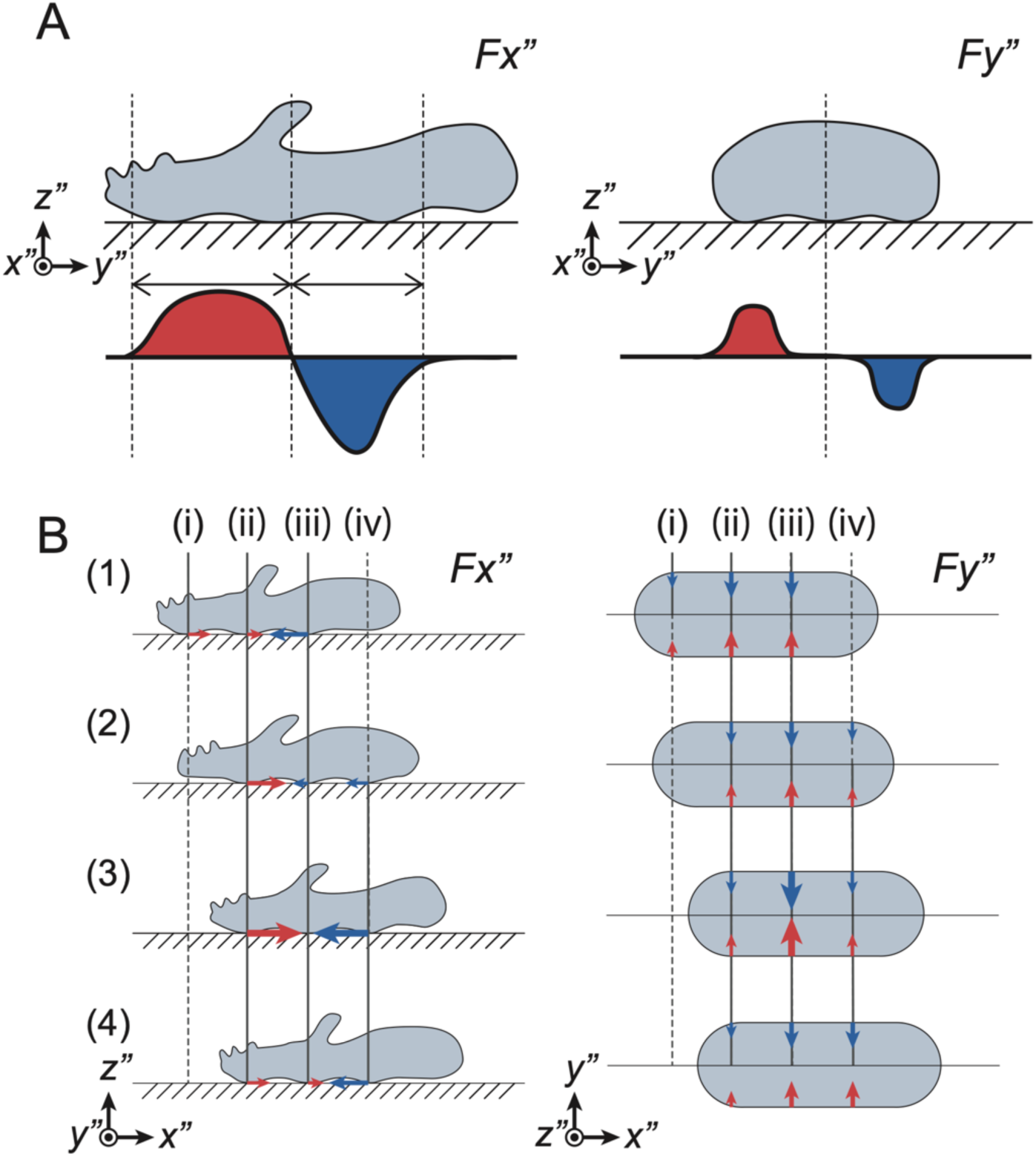
Schematics of traction stress patterns in *A. proteus*. (A) Schematic of the relationship between contact site position and traction stress in a cell. (Left) Component of traction stress along the migration direction. The upper schematic represents a cross-section parallel to the migration direction. (Right) Component of traction force perpendicular to the direction of cell migration. The upper schematic represents a cross-section perpendicular to the migration direction. (B) Schematic of the clawing pattern of *A. proteus*. (Left column) Motion of *A. proteus* and the distribution of the traction stress component along the migration direction, shown in a cross-section parallel to the migration direction. (Right column) Motion of *A. proteus* and the distribution of the traction stress component perpendicular to the migration direction, viewed from above. Arrows indicate the magnitude and direction of traction forces. Vertical dashed lines denote the positions where the cell adheres to the substrate. The dashed lines labeled (i)–(iv) indicate the positions where contact sites are formed. Solid segments along each dashed line represent the duration during which each contact site persists.

These observations also suggest that the periodic formation of contact sites at the front of the cell, followed by their detachment at the back, gives rise to a cyclic process of adhesion and traction stress generation that underlies amoeboid locomotion (Fig. 4B). In the following, we describe this cycle in more detail.

Focusing on the component of traction stress parallel to the migration direction (Fig. 4B, left column), newly formed contact sites at the leading edge (Fig. 4B, 2-iv) initially generate stress directed opposite to the migration direction. These stresses increase to a peak (Fig. 4B, 3-iv) and subsequently decrease (Fig. 4B, 4-iv and 1-iii). As these contact sites move toward the cell interior, the magnitude of the oppositely directed stress further decreases (Fig. 4B, 2-iii), approaches zero near the cell center (Fig. 4B, 3-iii), and then reverses direction to generate forward-directed stress as they move away from the center (Fig. 4B, 4-iii and 1-ii). With continued translocation, these contact sites (Fig. 4B, 4-iii and 1-ii) further increase forward-directed stress, reaching a peak (Fig. 4B, 3-ii), and then decrease in magnitude (Fig. 4B, 4-ii and 1-i). Finally, they detach from the substrate as they move further away from the cell center (Fig. 4B, 2-i). Importantly, contact sites at different stages of this cycle coexist, and their dynamics are coupled. Specifically, when a given contact site generates a strong backward-directed stress (Fig. 4B, 3-iv), an older site simultaneously reduces its stress magnitude (Fig. 4B, 3-iii), while an even older site generates strong forward-directed stress (Fig. 4B, 3-ii), which partially cancels the stress produced by more anterior sites. This coordinated sequence is repeated cyclically. Such a cyclic pattern in the parallel traction stress component was not observed in vegetative or poorly polarized *D. discoideum* undergoing random migration. In contrast, similar cyclic dynamics have been reported in chemotaxing *D. discoideum* and dHL60 cells[30]. These observations suggest that cyclic traction stress patterns are a general feature of highly polarized migrating cells.

Focusing on the component of traction stress perpendicular to the direction of migration (Fig. 4B, right column), contact sites formed at the leading edge gradually increased inward-directed lateral forces, reaching a maximum near the cell center. As these contact sites translocated toward the back of the cell, the magnitude of lateral stress decreased. Repetition of this process underlies A. proteus migration. A notable feature of this lateral stress component is that it peaks at the cell center.

This behavior differs from that of many other cell types, in which traction stresses are typically minimized near the center and exhibit a bipolar distribution with maxima at the front and back of the cell[29,30,39–41]. Quantitatively, the magnitude of the lateral traction stress was 0.85 ± 0.19 times (mean ± SD, N = 10 cells) that of the component parallel to migration. In contrast, in *D. discoideum*, which exhibits actin polymerization-driven amoeboid locomotion, this ratio is approximately 0.6[30]. These results indicate that *A. proteus* exhibits a relatively enhanced lateral component of traction stress. This distinctive stress distribution may be associated with the high motility of *A. proteus*.

It is worth noting that bleb-driven migration is frequently associated with an adhesion-independent regime characterized by extremely weak force transmission. Pioneering work has shown that certain fast-moving amoeboid cells, such as leukocytes, can bypass integrin-mediated attachments entirely, relying instead on shape deformation and cortical squeezing to navigate confined environments[24]. In such focal-adhesion-free modes, cells utilize nonspecific friction, and the propelling traction stresses are several orders of magnitude lower than those observed in classic mesenchymal migration, making them virtually challenging to resolve on standard two-dimensional TFM substrates[42]. In contrast, our findings demonstrate that *A. proteus* successfully generates robust, highly organized, and measurable 2D traction patterns despite its blebbing morphology.

This implies that *A. proteus* relies on a distinct mechanical regime where bleb formation coexists with efficient, localized substrate coupling. Our findings directly enrich the current classification of bleb-based migration strategies, highlighting that “blebology” encompasses a wider spectrum of mechanical regimes̶ranging from purely friction-based pushing to highly coordinated traction generation̶than previously appreciated[19]. Whether this characteristic spatiotemporal evolution of traction stress is a general feature of other bleb-driven amoeboid cells remains an open question.

In this study, we analyzed the relationship between amoeboid locomotion and the spatiotemporal characteristics of traction stress in *A. proteus*, a representative bleb-driven migrating cell. Our results show that *A. proteus* exhibits both traction stress features shared with actin polymerization-driven amoeboid cells and additional characteristics not observed in many other cell types. These results suggest that the mechanical mechanisms underlying amoeboid locomotion exhibit both universality and diversity.

## Methods

### a. Cell culture

*Amoeba proteus* was cultured in KCM medium (0.070 g/L KCl, 0.080 g/L CaCl₂, and 0.080 g/L MgSO₄·7H₂O) in 100 mm Petri dishes at 25°C in the dark. Cells were subcultured twice per week, and *Tetrahymena* (cultured in PPYG medium: 10 g/L proteose peptone, 2.0 g/L yeast extract, and 2.0 g/L D-glucose) was provided as a food source at the time of each subculture. One day after feeding, cells were collected from 100 mm dishes, washed with KCM medium, and transferred to 60 mm dishes filled with fresh KCM medium. Experiments were then performed on cells after an additional 24 hours of starvation.

### b. Measurement of traction stress

To prepare a disk-shaped polyacrylamide gel substrate, a silicone sheet (20 mm × 27 mm × 0.20 mm, Kenis Corporation) with an 8 mm circular hole punched at its center using a biopsy trephine was attached to a plasma-treated cover glass (24 mm × 32 mm, Matsunami). This assembly served as a mold for gel fabrication. The gel precursor solution, consisting of 4.4% (w/v) acrylamide, 0.20% (w/v) N,N’-methylenebisacrylamide, 0.14% (w/v) ammonium persulfate, 0.33% (v/v) N,N,N’,N’-tetramethylethylenediamine, and 0.24% (w/v) red fluorescent beads (1 µm carboxylated microspheres; Sicastar® RedF, Micromod), was poured into the mold.

The system was then covered with an untreated cover glass to form a sandwich structure and allowed to polymerize for 1 hour. After gelation, the upper plasma-treated cover glass was removed. The gel-attached cover glass was subsequently fitted with a second silicone sheet (17 mm × 25 mm × 3.0 mm, AS ONE) containing a rectangular opening (10 mm × 16 mm), such that the gel region was exposed within the opening. The chamber was then filled with KCM medium. The Young’s modulus of the gel was 9.615 ± 0.88 kPa (mean ± SD, N = 6), determined from the indentation depth (*Δd*) induced by placing a steel bead (diameter 1.0 mm, mass 4.0 ± 0.02 mg, mean ± SEM, N = 10) on the gel surface and analyzed using the Hertzian contact model[43].

For traction stress measurements, cells were placed on the gel and allowed to settle for 30 minutes before imaging cell motility and fluorescent bead positions. Imaging was performed using an inverted microscope (ECLIPSE Ti2, Nikon) equipped with a 20× Plan Apo objective lens (NA = 0.75, Nikon). Images were acquired using an ORCA-Fusion camera (Hamamatsu) with an exposure time of 50 ms. Bright-field images were obtained using a red-filtered LED light source (TI2-D-LHLED, Nikon), while fluorescence images were acquired using 550 nm excitation (D-LEDI-C, Nikon) and a TRITC filter set (Semrock). To track bead displacement on the gel surface, the focal plane was adjusted to the gel surface, and bright-field and fluorescence images were sequentially acquired. The beads used in this system were silica-based fluorescent microspheres localized at the gel surface. Imaging was performed at 5 s intervals. In addition, after the cell moved out of the field of view and the beads returned to their initial positions, the same imaging procedure was repeated, and these images were used as the reference initial bead configuration.

Fluorescence images were analyzed using particle tracking velocimetry (PTV) to determine bead displacements between frames. Bead detection and tracking were performed using the TrackMate plugin in Fiji[44,45]. Beads were detected using a Laplacian of Gaussian (LoG) detector with a detection radius of 0.33 µm and a threshold of 0, with subpixel localization enabled and no median filtering. Detected spots were filtered based on quality (QUALITY > 10). Tracking was performed using the sparse LAP tracker, with maximum linking and gap-closing distances of 0.98 µm. Splitting and merging events were not allowed. Tracks with a total displacement of less than 0.81 µm were excluded from further analysis. Prior to tracking, background subtraction was performed using a rolling-ball algorithm with a radius of 6.5 µm. To correct stage drift, translational motion was estimated from bead trajectories using TrackMate and subtracted on a frame-by-frame basis. To remove outliers corresponding to spatially localized, non-physical displacements, each bead was compared with neighboring beads within a radius of 15 µm. Beads whose displacement deviated from the local mean by more than one standard deviation were identified as outliers and excluded from further analysis. The resulting displacement images were divided into a square grid of 287 × 287 elements with a grid spacing of 2.6 µm. At each grid point, the mean displacement vector of beads located within a radius of 15 µm was assigned, yielding a two-dimensional displacement field (*u*_*x*_(*x*, *y*), *u*_*y*_(*x*, *y*)).

To estimate the stress field from the measured deformation field, we followed a previous study[46] and described the relationship between the substrate deformation field ***u***(***x***) and the stress field ***f***(***x***) using the integral form of the Boussinesq kernel ***G***:

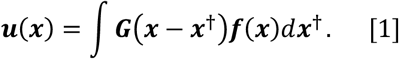

Here, *G* represents the integral Boussinesq Green’s function for a semi-infinite linear elastic substrate, obtained by integrating the classical point-force Boussinesq solution over a finite rectangular area element with uniformly distributed traction. For an incompressible substrate (*v* = 0.5), the kernel is given by

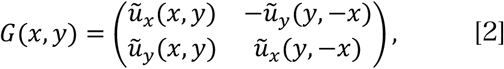

where

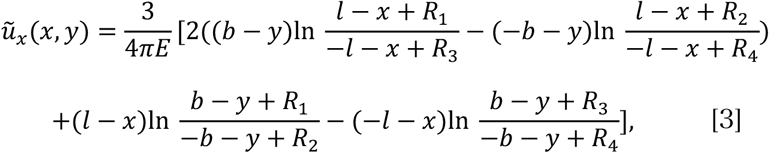

and

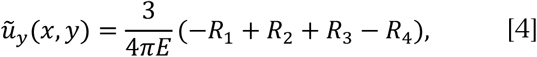

with

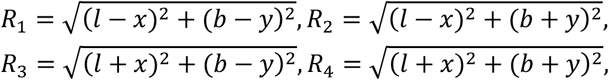

where *E* is the Young’s modulus of the substrate and 2*l* × 2*b* denotes the size of the rectangular force element (*l* = *b* = 2.2 µm). By applying a Fourier transform, Eq. [1] can be expressed in Fourier space as

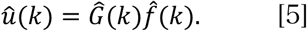

The stress field was then estimated by solving this inverse problem. The analysis was restricted to the cell region defined by a mask extracted from bright-field images, and the force outside the mask was set to zero. The cell mask was generated by first subtracting the background from the original image, computing the intensity gradient, and applying a Gaussian filter (σ= 3.25 µm) to the gradient image. The cell boundary was then extracted using the marching squares algorithm[47]. To reduce the influence of noise in the deformation field, the mask was expanded to approximately 1.2 times the original cell area. Deformations within this extended region were assumed to arise from forces generated within the original cell region and were therefore included in the subsequent analysis. This extended mask was chosen to ensure that most of the deformation field induced by the cell was captured. To obtain the optimal stress field, Tikhonov regularization[27,48] was applied by minimizing the functional

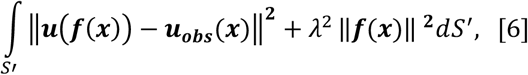

where the integration domain *S*′ corresponds to the extended cell mask. The symbol ∥⋅∥ denotes the Euclidean norm, and *λ* is the regularization parameter. Here, ***u_obs_***(***x***) represents the experimentally measured deformation field obtained from bead displacements, while ***u***(***f***(***x***)) denotes the deformation field predicted from the force field ***f***(***x***) via the forward model given by the Boussinesq kernel (Eq. [1]–[5]).

The value of *λ* was determined using the L-curve method[49] applied to randomly selected datasets, yielding a representative value of *λ* = 2.0 × 10^3^. The stress field ***f***(***x***) that minimizes Eq. [6] was obtained using an iterative optimization scheme based on the Jacobian.

### c. Characterization of traction stress

The cell centroid *x*_g_ was defined from the obtained force field as

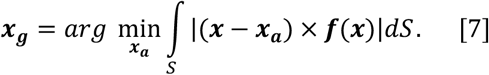

where *arg* min denotes the position that minimizes the integral quantity. is the traction stress vector, and ***x*** is the position vector. The integration domain *S*corresponds to the cell mask.

Next, we analyzed the temporal evolution of the traction stress distribution for each of the ten cells. At each time point, the cell centroid was used as the origin, and the cell velocity vector was temporally smoothed using a Gaussian filter (σ = 25 s) to define the direction of cell migration. A coordinate system was then defined such that the migration direction corresponded to the *x*″-axis and the perpendicular direction to the *y*″-axis. The traction stress field was transformed into this coordinate system, and was averaged over time and across samples to obtain a mean traction stress distribution.

Finally, for each time point, the spatial integrals of the traction stress components along the migration direction 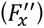 and perpendicular to it 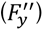 were computed, and their ratio was evaluated.

### d. Multipole analysis of traction stress

Multipole analysis was performed based on a previous study with minor modifications. The first moment (force dipole) was defined as

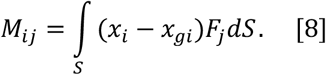

where the subscripts *i*and *j*denote Cartesian components taking values 1 and 2. *x*_1_and *x*_2_ correspond to the *x*- and *y*- coordinates, respectively, and *x*_g1_, *x*_g2_ denote the *x*- and *y*-components of the cell centroid. *F*_j_ represents the component of the traction stress vector. The resulting 2 × 2 tensor was diagonalized to obtain two eigenvalues and their corresponding eigenvectors. The eigenvector associated with the larger eigenvalue was defined as the major dipole, while that associated with the smaller eigenvalue was defined as the minor dipole. To quantify the relationship between the major dipole and cell migration, we calculated the distribution of angles between the major dipole and the cell velocity vector.

We further defined the second moment (quadrupole) as

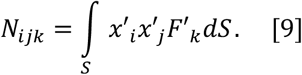

where the origin is fixed at the cell centroid, and the 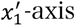 is defined such that it is most closely aligned with the direction of the major dipole. 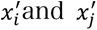 denote the position components in this coordinate system, and 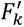 denotes the traction stress components. The force quadrupole tensor consists of six independent components: 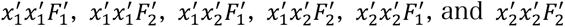. To investigate the relationship between these components and cell migration, we compared the sign of each quadrupole component with the sign of the velocity component along the *x*′-axis at each frame. The frequency of sign coincidence was defined as *P_sign_*. We further defined a correlation index *P* as

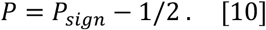

### e. Analysis of traction stress dynamics during cell motility

To analyze the spatio-temporal evolution of the traction stress distribution during cell migration, we constructed kymographs representing traction stress patterns along the direction of cell movement. Cross-correlations were then computed for different spatial shifts and temporal lags (Fig. 2E–G). In this analysis, the cell velocity vector was first temporally smoothed using a Gaussian filter (σ = 25 s). The cell centroid was then used as the origin, and a coordinate system was defined such that the direction of the smoothed velocity vector corresponded to the *x*″-axis, with the perpendicular direction defined as the *y*″-axis. Kymographs were constructed by decomposing the traction stress into *x*″- and *y*″-components (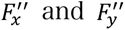) and integrating each component across the width of the cell, yielding *F*_*tx*_and *F*_*ty*_, respectively. For the *y*″-component, inward-directed forces were defined as positive to avoid cancellation between forces originating from the left and right sides of the cell.

Cross-correlation analyses were performed using the kymographs of *F*_*tx*_ and *F*_*ty*_, including correlations between *F*_*tx*_ (Fig. 2E), between *F*_*ty*_ (Fig. 2F), and between *F*_*tx*_ and *F*_*ty*_ (Fig. 2G). The cross-correlation function *C_AB_* (2D CCF) was defined as a root-mean-square–weighted Pearson correlation, with spatial shifts (axial position) Δ*x* and temporal lags *τ* as variables after normalization. The spatial shift Δ*x* was normalized by the average cell length *L*, and the temporal lag *τ* was normalized by the characteristic time *t*_0_, defined as the time required for a cell to travel its own length, *t*_0_ = *L*/*v*_0_, where *v*_0_ is the average migration speed.

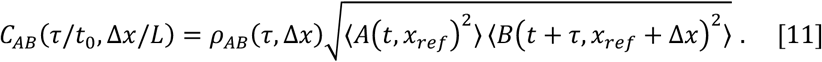

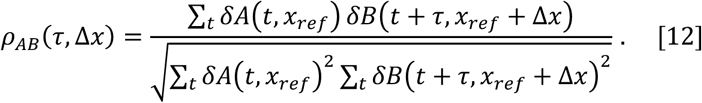

Here, ⟨⋅⟩ denotes temporal averaging, and *A* and *B* represent traction stress components (*A*, *B* ∈ {*F*_*tx*_, *F*_*ty*_}). The fluctuations are defined as δ*A* = *A* − ⟨*A*⟩ and δ*B* = *B* − ⟨*B*⟩. The reference axial positions *x*_ref_ for spatial shifts were determined by detecting peaks in the time-integrated traction-force magnitudes, ∑_*t*_ ∣ *F*_*tx*_ ∣ and ∑_*t*_ ∣ *F*_*ty*_ ∣, using a peak-detection algorithm[50] with a minimum peak height of 10 nN and a minimum peak-to-peak distance of 26 µm. The spatiotemporal cross-correlation function was calculated for each detected peak position, and the resulting correlation maps were averaged to obtain the representative correlation distribution. The resulting two-dimensional cross-correlation functions were averaged across 10 samples to obtain a mean cross-correlation map. Prior to averaging, each map was normalized such that its maximum absolute value was 1. To extract characteristic structures from the two-dimensional cross-correlation maps, peak detection was performed[47]. First, a Gaussian filter (σ = 2) was applied to reduce noise. Local extrema were then identified by comparing each point with its neighbors within a radius of 15 pixels (corresponding to a spatial shift of 0.15 and a temporal lag of 0.15). Points that were locally maximal or minimal were extracted as candidates. Finally, only points with an absolute correlation value greater than 0.1 were retained to identify significant peaks and troughs.

### f. Analysis of each contact site dynamics

To identify individual contact sites formed by cells, we first examined the magnitude of traction stress in each sample. Regions that did not exhibit clear and stable stress signatures consistent with contact site formation over the entire observation period were excluded from further analysis. The threshold for traction stress magnitude |***F***| was sample-dependent, ranging from 8 to 13 Pa. Subsequently, contact sites were defined as clusters of pixels exhibiting similar temporal dynamics in the traction stress field. Specifically, each pixel was characterized by the time series of the *x*-component (*F*_*x*_), *y*-component (*F*_*y*_), and magnitude (|***F***|) of the traction stress. Prior to clustering, these time-series data were smoothed using a three-dimensional Gaussian filter with standard deviations of one pixel (2.6 µm) in space and one frame (5 s) in time. Pixels were then clustered into two groups using the k-means algorithm[35,36], and clusters exhibiting coherent dynamics were extracted as candidate adhesion regions. For each pixel cluster, an elliptical fit was performed. Regions were retained as contact sites only if they satisfied the following criteria: area between 200–350 µm² or 1300–2500 µm², eccentricity ≤ 0.8–0.9, and solidity ≥ 0.8–0.9. Regions not satisfying these conditions were further processed: clusters with area ≤ 200 µm² were discarded, whereas larger regions were recursively subdivided. This procedure was repeated until only regions that satisfied all criteria remained (Fig. 3C).

To characterize the dynamics of each contact site, we analyzed both its position relative to the cell centroid and the traction stress generated within the region. Positions were defined in a coordinate system centered at the cell centroid, with the migration direction aligned with the *x*″-axis and the perpendicular direction defined as the *y*″-axis. The traction stress at each contact site was obtained by averaging the time-dependent traction stress across all pixels in the region. To visualize the spatial dependence of traction stress along the cell axis, contact sites were classified according to their position relative to the cell centerline. Specifically, positions along the *y*″-axis (perpendicular to migration) were divided into six bins ranging from -30 µm to 30 µm (bin width: 10 µm). For each *y*″ bin, the traction stress distributions along the cell axis *x*″ were averaged over all contact sites belonging to that bin. Only contact sites that could be tracked for more than 100 s and exhibited a positional standard deviation below 15 µm were included in the analysis.

Next, to quantify the spatial bias of the traction stress distribution, we calculated its stress-weighted axial position *x*″, defined as the weighted average of the coordinate *x*″ with the absolute traction stress as the weight:

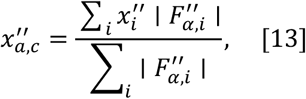

where 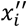 denotes the center coordinate of the i-th spatial bin, 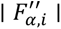 is the absolute value of the traction stress component in that bin, and *α* = *x* or *y*. This analysis was performed independently for each contact site and representative traction stress for each lateral position *y*″. For individual contact sites, the stress-weighted position was calculated only for contact sites whose mean occupied position along the *x*″-axis, defined as the average position of bins exhibiting nonzero traction stress, was within ∣ *x*″ ∣< 40 µm. This criterion ensured that only contact sites that could be tracked throughout their lifetime, from attachment to detachment, were included in the analysis. Representative traction stress distributions for each lateral position *y*″were obtained by averaging the normalized traction stress profiles of all contact sites belonging to the corresponding *y*″ bin, where each profile was normalized by its mean absolute traction stress.

## Data accessibility

Codes and data are available at https://doi.org/10.5281/zenodo.20652175.

## Declaration of AI use

The authors used ChatGPT (OpenAI) to assist with grammatical proofreading and stylistic improvements to the manuscript, and to develop code for some analyses and figures.

## Authors’ contributions

R. T.: conceptualization, data curation, formal analysis, investigation, methodology, software, validation, visualization, writing – original draft and writing – review & editing; S. E.: resources and writing – review & editing; C. F.: methodology and writing – review & editing; A. T.: resources and writing – review & editing; T. O.: validation and writing – review & editing; J. P. R.: methodology and writing – review & editing; K. S.: conceptualization, funding acquisition and writing – review & editing; T. N.: conceptualization, funding acquisition and writing – review & editing; Y. N.: conceptualization, methodology, resources, project administration funding acquisition, validation, supervision, writing – original draft and writing – review & editing.

## Supplementary Material

Supplementary Movie S1.

Bright-field movie of a migrating *A. proteus* (100× speed).

Supplementary Movie S2.

Time evolution of traction stress field during migration (100× speed). The white contour indicates the cell boundary. The color bar indicates stress magnitude. The red arrow denotes the major dipole, and the gray arrow indicates the cell velocity vector.

